# Virtual Clinical Trials of BMP4 Differentiation Therapy: Digital Twins to Aid Successful Glioblastoma Trial Design

**DOI:** 10.1101/2024.08.22.609156

**Authors:** Nicholas Harbour, Lee Curtin, Loizos Michaelides, Matthew E. Hubbard, Pamela Jackson, Vinitha Rani, Rajappa S. Kenchappa, Virginea de Araujo Farias, Anna Carrano, Markus R. Owen, Alfredo Quinones-Hinojosa, Kristin R. Swanson

## Abstract

Glioma stem cells (GSCs) are considered a major driver of glioblastoma (GBM) progression and are highly resistant to standard cytotoxic treatments. BMP4 has been shown to drive differentiation of GSCs, increase sensitivity to radiotherapy, slow growth and increase survival times in animal models. To assess the potential of BMP4 as a differentiation therapy, we develop a mathematical model that describes the growth of a GBM tumor via a hierarchy of GSCs, progenitor cells and terminally differentiated cells. We parametrize our model using experimental data from twelve patient-derived GSC lines, on which we measured response to radiotherapy and population growth with and without exposure to BMP4. Cell lines were typically more sensitive to radiotherapy after two days of BMP4 treatment but population growth can either increase or decrease after seven days of exposure to BMP4. To identify key parameters that drive successful treatment we perform global sensitivity analysis which identifies key parameters for BMP4 efficacy including proliferation rate and self-renewal sensitivity of GSCs. We then compare two treatment schedules: a single dose of BMP4 at resection and continuous delivery of BMP4 from resection till the end of radiotherapy. Due to the short half-life of BMP4 and its synergy with radiotherapy, continuous delivery of BMP4 during radiotherapy is more effective than a single dose prior to radiotherapy. We then perform a series of virtual clinical trials, stratified by tumor proliferation rate and GSC self-renewal sensitivity, which allows us to estimate the probability of observing a successful early-phase clinical trial for various virtual patient cohorts. We find that trials that selected the subset of patients with more proliferative GBMs were more likely to lead to significant improvements in survival.

**Significance:** Targeting glioma stem cells with BMP4 provides a novel opportunity to shift the complex cellular ecosystem of gliomas to enhance treatment efficacy. Mathematical modelling can facilitate optimal patient tumor feature selection when designing successful clinical trials.

## 1. Introduction

Glioblastoma (GBM) is the most common primary malignant brain cancer in adults (1,2). Current standard of care consists of maximal safe surgical resection followed by concurrent radiotherapy and chemotherapy (with temozolomide) (3). Despite this aggressive treatment and repeat surgical resections (4), outcomes remain poor with five-year survival rates of only around 7% (1,2). The aggressiveness and fatal outcomes of GBM can be attributed in part to the presence of a small population of cells with stem-like characteristics, known as glioma stem cells (GSCs) (5–13). GSCs are a major driver of tumor progression that can initiate brain tumors in mouse models of GBM (8). In addition to being tumor-initiating, GSCs are highly resistant to both radio- and chemo-therapy due to activation of highly efficient DNA repair mechanism (10,12,14–18). Consistent with this, studies have found that the percentage of GSCs is increased after radiation exposure (14,19). Worryingly, experimental and simulation results show that the surviving GSCs are able to initiate tumor recurrence with enhanced aggressiveness and decreased latencies relative to untreated tumors (14,20).

Evidence for a proliferative hierarchy in GBM has been observed in many experiments including in xenotransplantation (7,8), lineage tracing in mouse models (21), *in vitro* genetic barcoding (13) and single-cell RNA sequencing (22,23). In all these cases it is proposed that slow-cycling stem-like cells give rise to a more rapidly cycling progenitor-like population of cells (PCs) with extensive but finite self-maintenance capacity, that ultimately generates terminally differentiated non-dividing cells (TDCs). The identification of GSCs as a restricted subpopulation of cells responsible for the maintenance of tumors suggests that these are the cells that have the inherent ability to drive cancer evolution and seed recurrence. Therefore, if treatment outcomes are to be improved for patients with GBM, it is vital that therapeutics that specifically target GSCs are developed (15,24,25).

One potential approach to target GSCs is differentiation therapy. This is based on the idea that signals promoting differentiation could be effective at driving malignant stem-like cells to a less aggressive and ideally post-mitotic differentiated state (26–28). This approach has seen success in acute promyelocytic leukemia (APL) where all-trans-retinoic acid (ATRA) promotes differentiation of cancer stem cells and can lead to complete remission (29,30). In GBM, bone morphogenetic protein 4 (BMP4), a member of the TGF-*β* superfamily, has shown potential as a differentiation therapeutic agent. BMP4 has been shown to drive differentiation of GSCs towards a predominantly glial (astrocytic) fate, to reduce GBM tumor burden *in vivo* and to improve survival in a mouse model of GBM (26,31–34). Additionally, studies from our group and others have shown that adipose-derived mesenchymal stem cells (AMSCs) provide a possible delivery mechanism, as they can be non-virally engineered to secrete BMP4 (or other therapeutics) (33,35,36). These AMSCs preferentially migrate toward areas of malignancy, homing in on infiltrating glioma cells, gradually releasing their cargo of BMP4 (32). Furthermore, BMP4 recently (2023) completed its first in-human phase I clinical trial (ClinicaTrials.gov identifier: NCT02869243) (37). In this study, BMP4 was delivered locally via convection-enhanced delivery (CED) and was found to be safe and well tolerated, with no severe drug-related adverse events. While efficacy was not the primary endpoint, encouraging signs were observed: of fifteen patients treated (all recurrent GBM), three (20%) responded, including two who achieved complete responses with long-term survival. These results provide compelling support for pro-differentiation therapy as a viable treatment avenue in GBM and strongly motivate further investigation to better evaluate the therapeutic potential of BMP4.

Even with the most promising preclinical data, designing clinical trials for novel therapies is fraught with challenges, including patient selection criteria, intra-tumor heterogeneity, and variability between patient populations. Clinical trials typically consist of four phases in which a new intervention is investigated in human subjects to determine its safety and efficacy. This system is notoriously resource-intensive and inefficient. The average cost per patient is $59,500 and takes more than 10 years, with only 10% of drugs in phase I studies eventually approved (38). In GBM, the statistics are even worse; multiple recent phase 3 trials have failed to meet their prespecified primary endpoints (39). Standard of care as well as survival rates have remained largely unchanged since the introduction of the Stupp protocol in 2005 (3,40). Of course, there are many reasons for these failures including inter- and intra-patient variability, drug delivery limitations, paucity of control arms, overly stringent clinical eligibility, and beyond (38,39,41). Digital twins simulating virtual tumors and their therapeutic response provide a promising new strategy to reduce the need for animal and human testing while streamlining the development of novel therapies that can ultimately be tested in human clinical trials (42,43). The digital twin is a system of mathematical equations that encode key prior biological knowledge of how cells interact during tumor growth and in response to therapy (44–46). Solving these equations allows us to simulate the tumor over time, enabling comprehensive analysis of the dynamics over the duration of disease in a manner that is not possible in humans or any current biological modelling framework (46). By thoroughly testing the model first *in silico*, simulating virtual clinical trials, we can gain insight into the potential outcomes of a clinical trial before it is conducted, allowing us to maximize the chance of success and minimize wasted resources (47,48).

In this article we develop a digital twin for GBM tumors based on an ordinary differential equation (ODE) model that simulates the structured hierarchical growth of GSCs, PCs and TDCs as well as response to radiotherapy and differentiation therapy. While we do not claim to capture the full complexity of a GBM tumor, we believe that our model captures the key non-linear dynamics that are critical for characterizing the effects of any potential differentiation therapy such as BMP4. The rest of this article is structured as follows: In the materials and methods section we present the *in vitro* cell line data used to develop and parametrize the model, the mathematical model itself (including how we simulate differentiation therapy and radiotherapy) and how we parametrized the model by simulating the *in vitro* experiments. In the results section, we show that our model can reproduce the experimentally observed dual outcomes of BMP4 exposure *in vitro*, where some cell lines grow more and others grow less after seven days of BMP4 exposure, when compared to the control without BMP4. We performed global sensitivity analysis (GSA) on the model to identify the key parameters that drive successful treatment as well as compare two potential treatment schedules: a single dose of BMP4 at resection and continuous delivery of BMP4 from resection to the end of radiotherapy. Finally, we performed a series of virtual clinical trials to predict the probability of observing a successful early-phase trial for a series of stratified patient cohorts based on the important parameters identified in the GSA. The paper workflow is summarized in Figure 1.

**Figure 1:**
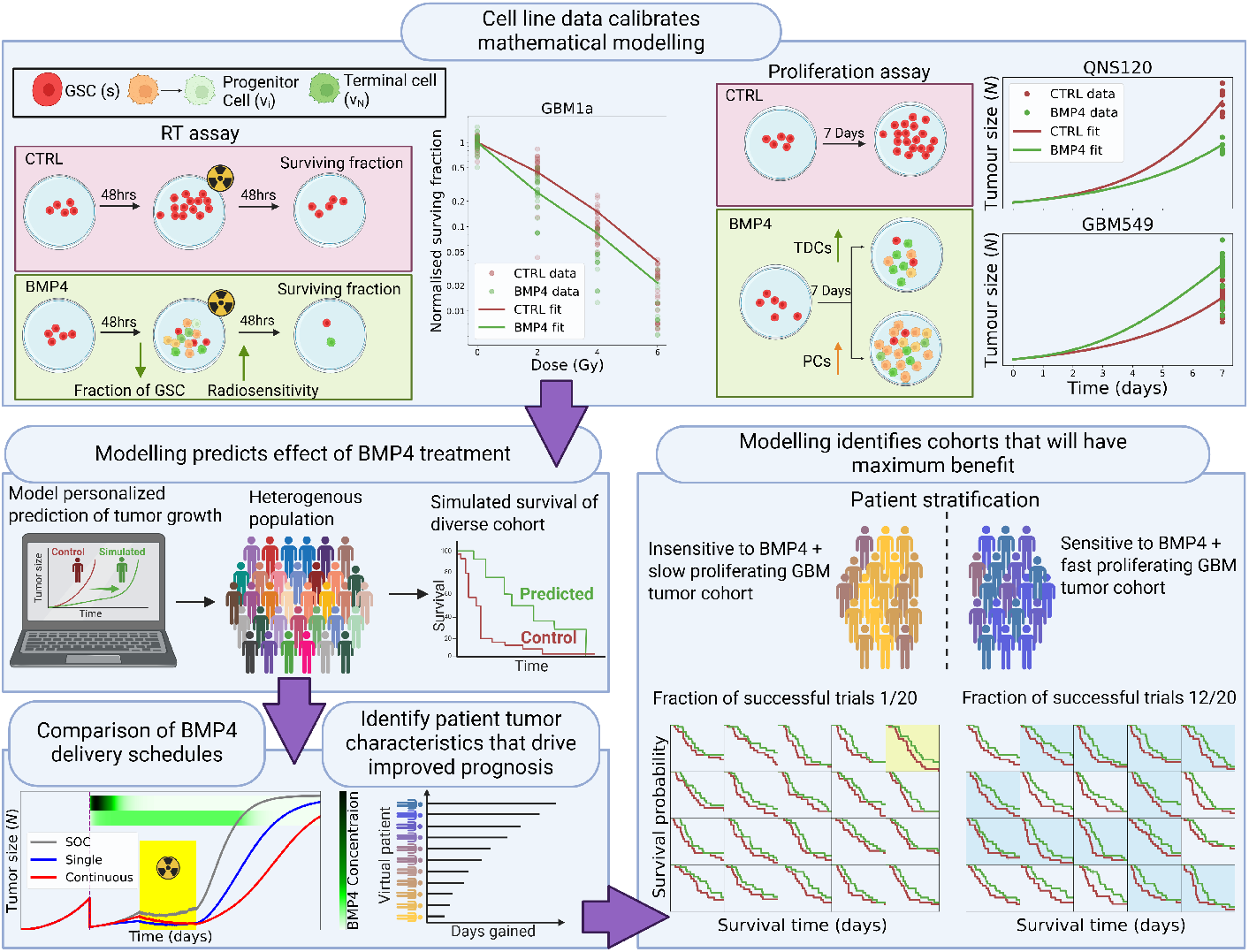
Digital twin virtual clinical trial workflow. Experimental data from *in vitro* experiments feeds into the development and calibration of the mathematical model that forms the digital twin. Using our model, we can predict the effect of BMP4 therapy on a diverse heterogenous population. We can compare and optimize different delivery schedules for BMP4 therapy. This allows us to identify patient tumor characteristics that result in improved survival. Based on these characteristics we can predict patient cohorts that will derive maximum benefit from BMP4 treatment that should be selected for in clinical trials. (created with BioRender.com)

## 2. Materials and methods

### 2.1. Cell line and culture conditions

The GBM cell lines were derived from adult glioblastoma patients using our previously established protocols (49). The cell lines were sub-cultured in DMEM/HAM-F12 media supplemented with 1% antibiotics, 1% glutamine and 2% Gem 21 neuroPlex supplement and maintained at 37 ºC at 5% CO2 in a sterile incubator throughout the study. GSC doubling time, as well as metadata including sex and primary/recurrent status are provided in Table 1.

**Table 1:** Fitted parameter values and metadata for all patient-derived GSC cell lines. Cell lines are order by doubling time.

| Cell line | Primary/<br>Recurrent | Sex | Doubling time<br>(hrs) | Radiosensitivity<br>($\alpha$) | GSC self-renewal<br>sensitivity ($\psi$) | Compartment<br>sensitivity ($\varphi$) |
| --- | --- | --- | --- | --- | --- | --- |
| GBM965 | P | F | 36.1 | 0.134 | 0.002 89 | 0.0789 |
| QNS120 | P | M | 43.5 | 0.116 | 0.001 59 | 0.0975 |
| GBM522b | R | M | 53.4 | 0.199 | 0 | 0 |
| GBM1a | P | M | 54.7 | 0.338 | 0.0108 | 0.0737 |
| GBM626 | P | M | 57.1 | 0.328 | 0.101 | 0.0635 |
| GBM549 | P | F | 60.4 | 0.189 | 0.007 15 | 0.0494 |
| QNS315 | P | F | 63.6 | 0.0841 | 0 | 0 |
| QNS657 | P | F | 75.6 | 0.104 | 0.0245 | 0.0723 |
| QNS679 | R | F | 108 | 0.146 | 0.197 | 0.0614 |
| GBM640 | P | F | 109 | 0.108 | 0.192 | 0.0864 |
| QNS108 | P | M | 109 | 0.151 | 0.003 11 | 0.0782 |
| QNS166 | P | F | 117 | 0.0872 | 0.0116 | 0.0751 |

### 2.2. Radiotherapy assay

Cells were treated with stem cell media (untreated control) and BMP4 conditioned media (100 ng/ mL), for a period of 48 hours. Then, seeded at the rate of 250-1250 cells per well in sextuplicate in 96 well plates and treated with 0, 2, 4, 6 Gy of radiation. After 14 days the colonies with more than 100 nm diameter were manually counted. The results are plotted in Figure 2 (A).

**Figure 2:**
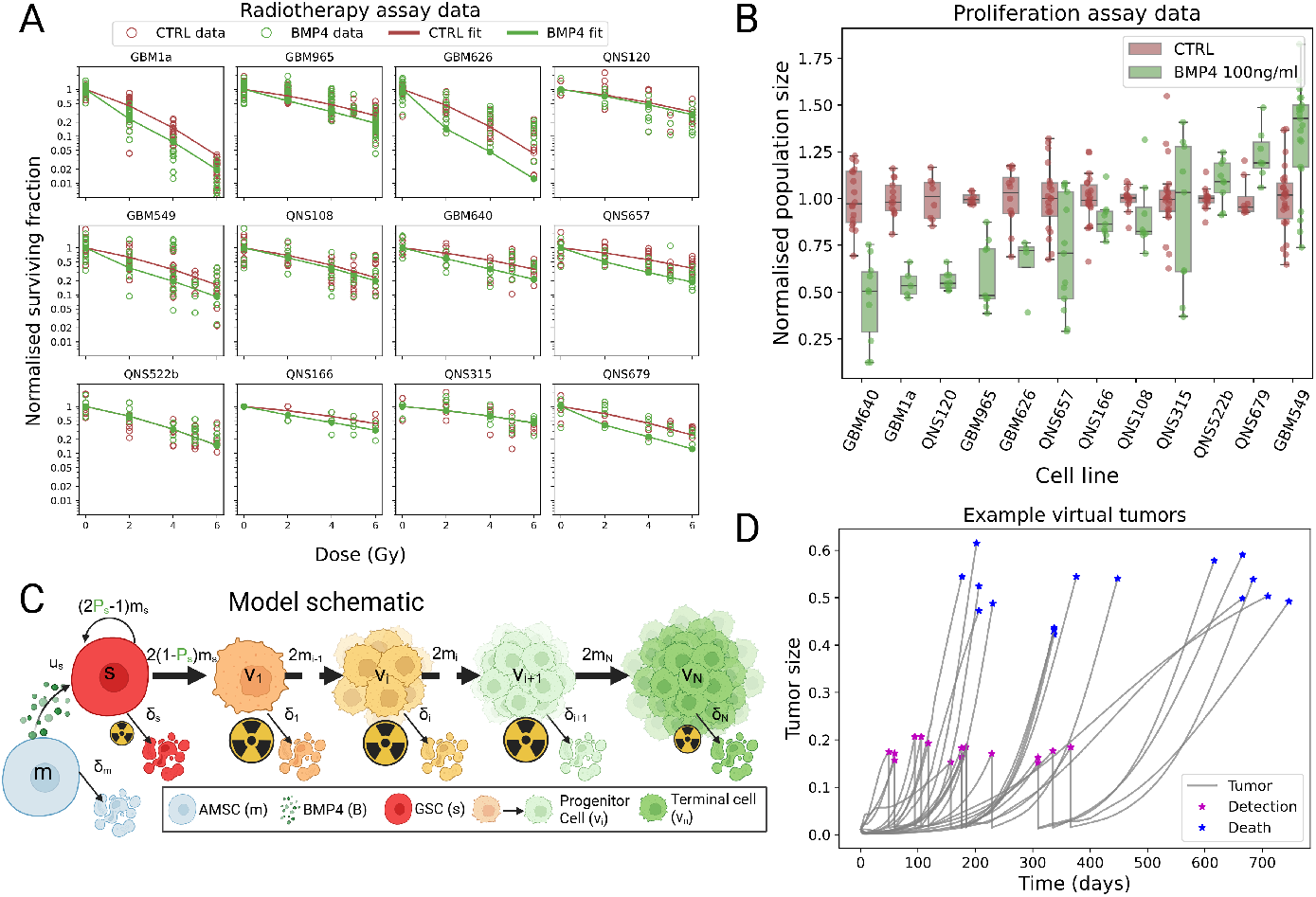
Cell line data. A) Data from radiotherapy assay on all twelve patient-derived GSC cell lines. Raw data points shown in circles and the model fit shown with solid lines. In general BMP4 increased sensitivity to radiotherapy. B) Data from the proliferation assay. Majority of cell lines showed a decreased total size in the BMP4 case, however four cell lines showed an increase in average size after BMP4 exposure. C) Our mathematical model is based on this schematic for GSC proliferation, GSCs are at the base of a hierarchy of cellular division ultimately ending in terminally differentiated cells (created with BioRender.com). D) Examples of total tumor size for 20 virtual patients. All virtual patients start with the same size initial condition and growth diverges due to difference in parameters and stochasticity in detection times. Once the tumors have grown to size of around 0.2 they are detected and surgery, followed by RT, is simulated. Ultimately the simulation is randomly ended, simulating a random death event that increase in likelihood as the tumor grows larger, typically occurring once the tumor has reached a size around 0.5-0.6.

### 2.3. Proliferation assay

GSCs were seeded at 1,000 cells/well in sextuplicate in 96-well plates and treated with 100 ng/mL of BMP4 for a period of 7 days. We quantified the cell proliferation using Cyquant cell proliferation assay kit (C7026; Invitrogen, USA) as described by manufacturer’s instructions. The fluorescence was measured using an HTX Synergy microplate reader (BioTek Instruments, Inc; Winooski, VT, USA) using an excitation wavelength of 480 nm and an emission wavelength of 520 nm. The results are shown in Figure 2 (B).

### 2.4. Virtual tumor model

To simulate a virtual GBM tumor we assume that it is comprised of a hierarchy of GSCs and PCs with decreasing proliferative capacity, ultimately leading to TDCs. The main assumption on GSCs, PCs and TDCs are:

- GSCs have unlimited replicative potential.
- PCs have a limited replicative potential (*n*), which ensures that they are never tumor-initiating. Once PCs have reached compartment *n* they become TDCs and can no longer proliferate.
- GSCs are capable of both self-renewing and differentiating. We denote the proportion of GSCs that self-renew by *P*_*s*_. This does not distinguish between symmetric and asymmetric division, but rather just considers the overall fraction of GSCs that self-renew (*P*_*s*_) and differentiate (1 − *P*_*s*_). Note that it can be shown that alternative formulations that specifically account for asymmetric, symmetric differentiation and symmetric self-renewal of GSCs are equivalent to this (20,50).
- In general, we allow GSCs to differentiate into PCs with any proliferative capacity, the fraction of differentiation into each compartment is given by the vector ***r***. We assume that without BMP4, differentiation is perfectly hierarchical, that is all GSCs differentiate into PCs in compartment *i* = 1 and divide exactly *n* times before becoming TDC, which corresponds to *r*_1_ = 1 and *r*_*i*_ = 0 for *i* = 2, …, *n*.

Figure 2 (C) shows a schematic of this model. Following these assumptions, we derive the following system of equations:

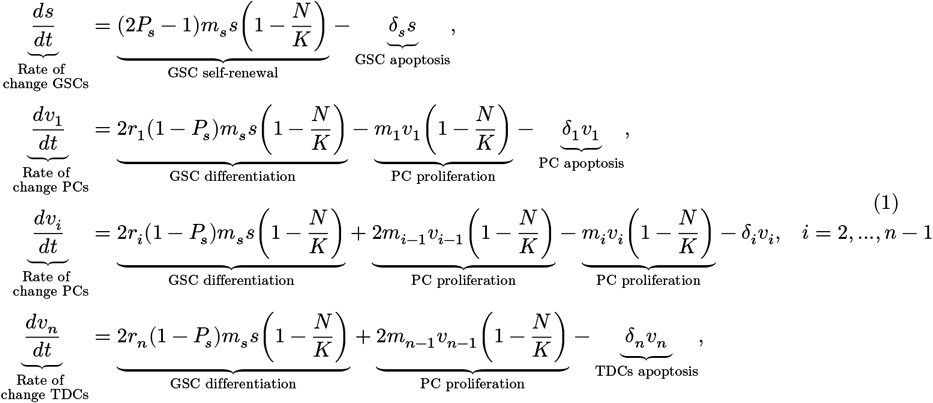

where *s*(*t*) is the density of GSCs, *v*_*i*_(*t*) is the density of PCs (in compartment *i*) and *v*_*n*_(*t*) are TDCs. The total size of the tumor is given by 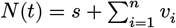. The apoptosis rates are given by *δ*_s_ for GSCs and *δ*_*i*_ for PCs. The proliferation rates of GSCs and PCs are given by *m*_*s*_, *m*_*i*_ respectively, where for simplicity we assume all *m*_*i*_ are equal. These proliferation rates are subject to a crowding response, represented by the term 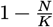 where *K* is the carrying capacity of the tumor. Throughout, following (19) we assume that PCs have a maximum proliferative capacity of *n* = 10.

### 2.5. BMP4 delivery model

We model the delivery of BMP4 via adipose-derived mesenchymal stem cells (AMSCs). The AMSCs simply decay exponentially from an initial concentration at implantation (33). BMP4 is released from these AMSCs and taken up by GSCs. From these assumptions, we derive the following system of equations:

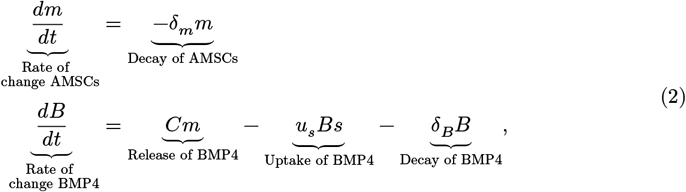

where *m* is the density of AMSCs and *B* is the density of BMP4. *δ*_*m*_ is the decay rate of AMSCs, *C* is the rate at which AMSCs release BMP4, *u*_*s*_ is the uptake rate of BMP4 by GSCs and *δ*_*B*_ is the decay rate of BMP4. *In vivo* experiments have shown that AMSCs can survive for around 14 days in a rodent model and that BMP4 reaches its peak concentration at around 48hrs after initial implantation of AMSCs (33). Throughout this article, we will refer to BMP4-AMSC delivery simply as BMP4, since the two are functionally equivalent apart from a short delivery delay.

### 2.6. Differentiation therapy model

Following (28), we model differentiation therapy through a simple relationship between the level of differentiation promoter (BMP4), and the probability of GSC self-renewal, *P*_*s*_, given by

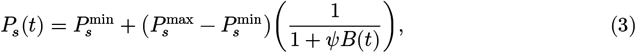

where 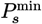 and 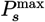 are the minimum and maximum self-renewal probabilities and *B*(*t*) is the concentration of BMP4. Self-renewal is maximal, 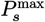, when there is no BMP4 present, and decreases to a minimum value 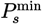 as BMP4 increases. We do not consider any endogenous production of BMP4 (or any other differentiation promoter), therefore it is only present during prescribed differentiation therapy. The parameter *ψ* represents the GSC self-renewal sensitivity to BMP4, as *ψ* increases (for a fixed concentration of BMP4) 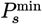 is approached faster, see Figure 3 (A).

**Figure 3:**
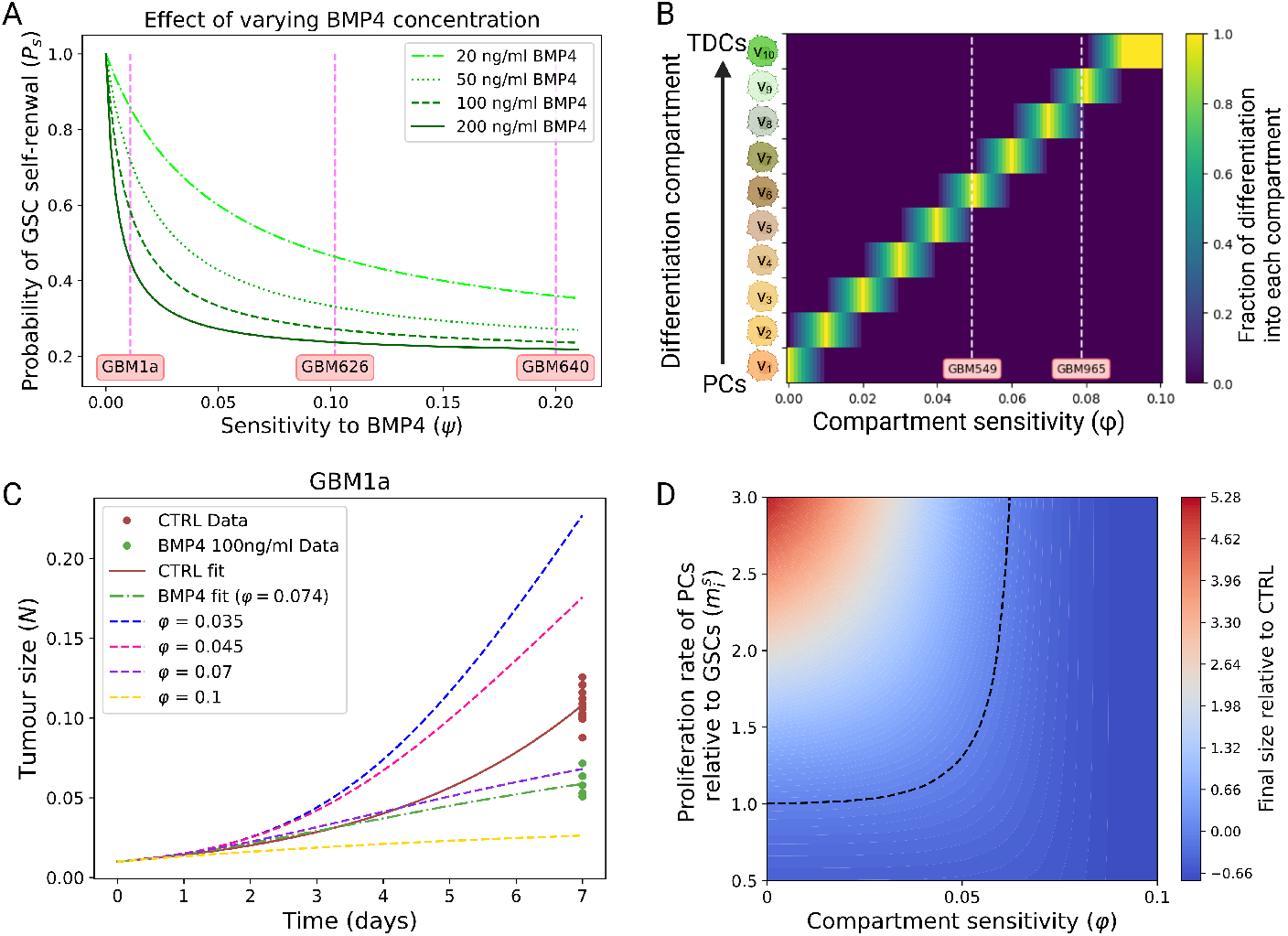
A) Probability of GSC self-renewal depends on concentration of BMP4 and cell line sensitivity to BMP4. As sensitivity is increased *P*_*s*_ decreases according to Equation 3; this relationship is shown for various fixed concentrations of BMP4. B) Compartment sensitivity (*φ*) dictates the proliferative capacity of progenitor cells according to Equation 5 (created with BioRender.com). C) Simulation of the proliferation assay with parameters taken to match cell line GBM1a and *φ* varying. D) The key parameters that influence the final size after seven days are the compartment sensitivity (*φ*) and the relative proliferation rate of PCs to GSCs, 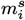. Taking parameters from the GBM1a cell line, we plot a heat map of the final size (relative to CTRL); the black dashed curve indicates where both CTRL and BMP4 arms are the same size after seven days.

In addition, we also assume that BMP4 can alter the proliferative capacity of the resulting PCs. That is, in the presence of BMP4 a GSC that differentiates into a PC may only divide a small number of times or go straight to a TDC (26,34). We implement this in our model as follows. Let *i* ∈{1, 2, …, *n*} index the compartment of the PCs. We define the piecewise linear triangle function centered at zero

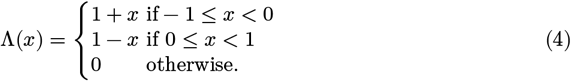

Let *θ* = *φB*(*t*) be the center of the triangle response. Then the PC differentiation state distribution ***r***, i.e., the proportion of GSC cell divisions into compartment *i*, is given by

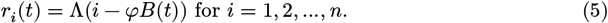

This form means that as compartment sensitivity *φ* gets larger (or BMP4 concentration), PCs produced from GSC division have reduced proliferation capacity. For the most sensitive cell lines this will mean that GSCs differentiate directly into TDCs. How ***r*** changes with *φ* and BMP4 concentration is visualized in Figure 3 (B).

### 2.7. Radiotherapy model

To model the effects of RT we use the linear quadratic (LQ) model, widely used in mathematical modelling of RT (51–54), but modify it to account for the different sensitivities to RT between GSCs, PCs and TDCs. Therefore, the fraction of cells that survive, *γ*_rad_(*d*), after a single fractional dose of *d* Grays (Gy) of radiation is given by

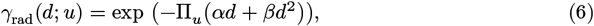

where *α* can be interpreted as death from single-strand breaks (linear component) and *β* can be interpreted as death from double-strand breaks (quadratic component). Throughout, in order to reduce the number of parameters we fix *β* = *α*/10 as has been done previously (51–53). The term Π_*u*_ accounts for the differential radiosensitivity of the cell subpopulations in the model. GSCs are less sensitive to radiation than other cancer cells, thus we introduce the additional radio-protection parameter *η* with Π_*s*_ = *η*. Previous studies have estimated GSC radio-protection to be *η* = 0.1376 (19). RT is also less effective on terminally differentiated cells, so we include an additional parameter *μ*, with 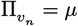, to account for the radio-protection of non-proliferating cells. Previous studies have estimated this to be *μ* = 0.5 (19). We assume that all stages of progenitor cells are equally sensitives to RT and thus 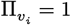 for *i* = 1, 2, …, *n* − 1 It is assumed that all effects of radiation on tumor cell density are instantaneous, and no delay or otherwise toxic effects of radiotherapy are considered. Nor are any effects on proliferation rate as a result of radiotherapy considered. Thus, we model the effects of radiotherapy as an instantaneous reduction of tumor density, applied to each compartment in the model separately.

### 2.8. Resection model

We model resection as an instantaneous loss of density applied equally to GSCs, PCs and TDCs. Previous studies have estimated that on average resection results in a 91.7% reduction in tumor volume (55). Accordingly, in our model resection is implemented as an instantaneous 91.7% reduction distributed equally across all cellular subpopulations.

### 2.9. Parametrizing sensitivity to BMP4

To parametrize the key model parameters of BMP4 sensitivity *ψ* and *φ* we simulate both the RT assay and the proliferation assay, jointly minimizing the overall error between the simulated and experimental data across both simulated assays. An example of the results of this parameter fitting is given in Figure 1. For the corresponding plots across all cell lines see supplementary Figure S1. In all simulations we use the measured cell line doubling times to set the GSC proliferation rate *m*_*s*_, and the other parameters used are fixed across all cell lines according to the values in Supplementary Table S1. The final fitted parameters for each cell line are shown in Table 1.

### 2.10. Virtual patient simulation

To simulate a realistic virtual patient’s progression, we incorporate stochasticity into the tumor detection and survival times. We assume that both detection of the tumor and death depend on tumor size, subject to random variation, for each virtual patient. Detection (death) is more likely the bigger a tumor is, but two tumors of equal size in two patients do not necessarily lead to detection (death) at exactly the same times. Thus, we assume that the times of tumor detection and death, represented by the random variables *T*_detect_ and *T*_death_, depend on total tumor density *N*(*t*) according to the hazard function

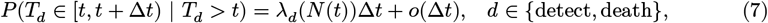

where

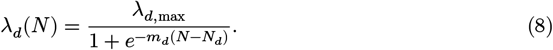

Here, *λ*_*d*(*N*)_ is a shifted logistic function; *λ*_*d*,max_ is the maximum rate of detection (which we take to be 1), *m*_*d*_ describes the steepness of the logistic function (which we set to be 100 in the detection case and 20 in the death case). The constant *N*_*d*_ is a threshold parameter at which the probability rate of detection or death is half-maximal (for each of *d* ∈ {detect, death}), which we set to be 0.2 and 0.7 in the detection and death cases, respectively. Δ*t* is the time step on which the model is solved numerically. This is similar to the approach of (56,57), apart from our choice of nonlinear dependence on tumor size. The shifted logistic function acts as a switch mechanism, meaning once *N* > *N*_*d*_ the rate of detection (death) rapidly increases towards its maximum. This leads to detection and death times that are random but clustered around specific sizes of tumor, see Figure 2 (D).

Additionally, to simulate realistic virtual patients, we also must simulate standard of care treatments. Once the tumor is detected we simulate immediate resection, then after thirty days RT is simulated, according to Equation 6, in a typical five days on two days off schedule where each day 2 Gy of radiation is delivered over the course of six weeks for a total RT dose of 60 Gy (3).

### 2.11. Data availability

All data for the cell lines as well as code for parameter fitting, model simulation and virtual trials, required to reproduce all the work presented in this article are available at: https://github.com/Harbour-N/Virtual-Clinical-Trials-of-BMP4-Differentiation-Therapy.

## 3. Results

### 3.1. BMP4 increases sensitivity to radiotherapy in a subset of cases

Figure 2 (A) shows that two days exposure with BMP4 prior to radiotherapy increases sensitivity in several patient-derived cell lines, to varying degrees. To design and implement clinical trials of BMP4 therapy, it is essential to obtain quantitative measures of each cell line’s sensitivity that could help guide patient selection and stratification.

We believe that the variability in radiotherapy response after BMP4 exposure arises from changes in cellular subpopulations driven by BMP4-induced differentiation. This effect is largely mediated by the GSC self-renewal sensitivity, *ψ*, which quantifies how the probability of GSC self-renewal changes in response to BMP4 (Equation 3). The resulting cellular composition after two days is then determined by a combination of two key factors: the proliferation rate (doubling time) of the cell line and the probability of GSC self-renewal. For example, in a fast-dividing cell line, a modest decrease in *P*_*s*_ can produce the same GSC-to-non-GSC ratio as a slower-dividing cell line in which BMP4 induces a stronger reduction in self-renewal. Consequently, simple comparisons of relative survival between cell lines is insufficient; a mechanistic model is needed to disentangle these potentially confounding effects.

This distinction is illustrated by comparing GBM1a and GBM640 in Figure 2 (A). At 4 Gy in the CTRL condition, mean survival is 0.1163 for GBM1a and 0.4167 for GBM640. Following BMP4 treatment, survival falls to 0.044 in GBM1a and 0.267 in GBM640. This corresponds to a 2.6-fold reduction in GBM1a versus a 1.5-fold reduction in GBM640. However, because GBM1a proliferates more rapidly, the model predicts that BMP4 actually exerts a stronger effect on reducing GSC self-renewal in GBM640 than in GBM1a. The fitted sensitivities and doubling times for all cell lines are reported in Table 1.

### 3.2. Dual outcomes of differentiation therapy in GBM cell lines

Data from the proliferation assay shows that compared to control (CTRL), in the majority of cell lines, exposure to BMP4 leads to a reduced size after 7 days, but in some cases it can lead to an increase in size after the same time period (Figure 2 (B)). To investigate this dual behavior we used our model to simulate the proliferation assay experiment. As in the real *in vitro* assay, we initialized simulations with a small number of GSCs and consider a CTRL and BMP4 arm. In the CTRL experiment (stem cell media), we assume that no differentiation takes place, i.e., *P*_*s*_ = 1, while in the BMP4 arm we assume a constant concentration of 100 ng/ml of BMP4 is present for the 7 days. The doubling times, along with the fitted self-renewal sensitivity (*ψ*) and compartment sensitivity (*φ*) are given in Table 1. Figure 1 shows the results of these simulations for two specific cell lines QNS120, which decreases in size, and GBM549, which increases in size. The equivalent plots for the other cell lines are shown in Supplementary Figure S1.

Figure 3 (B) illustrates what different compartment sensitivities *φ* represent in terms of the proliferative capacity of PCs. In particular, we show how the vector ***r*** (Equation 5), which represents the proportion of GSC divisions into each PC compartment, changes as we vary *φ*, for a fixed concentration of BMP4 of 100 ng/ml. In the case of GBM549 the fitted compartment sensitivity is *φ* = 0.0494, which corresponds to the majority of GSCs differentiating into compartment six (*v*_6_) - or equivalently PCs that can proliferate four more times before becoming TDCs. While for GBM965 the fitted compartment sensitivity is *φ* = 0.0787, which corresponds to the majority of GSCs differentiating into compartment nine (*v*_9_) - or equivalently PCs that proliferate once before becoming TDCs. In Figure 3 (C), we show how varying the compartment sensitivity affects the results of the proliferation assay. Growth rate and sensitivity parameters are chosen to match cell line GBM1a. This shows how varying the compartment sensitivity can give rise to the observed dichotomy in the data. Enhanced differentiation can potentially give rise to more growth as PCs are assumed to proliferate faster than GSCs. While on the other hand if PCs are pushed too far towards TDCs (increasing *φ*) it can lead to a decrease in size due to fewer proliferating cells.

The other key parameter in determining the size after seven days of the CTRL and BMP4 cases is the difference in proliferation rate between GSCs and PCs, 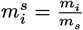. Thus far, for simplicity, we have assumed that PCs proliferate twice as fast as GSCs. Figure 3 (D) shows the difference in size (BMP4 size / CTRL size) after seven days of BMP4 exposure for various pairs of compartment sensitivity (*φ*) and 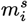, taking all other parameters fixed to match the GBM1a cell line. The black dashed line indicates the contour at which the CTRL and BMP4 arms are the same size at seven days. If PCs proliferate more slowly than GSCs, that is 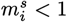, then as would be expected, we always observe a smaller tumor cell population after seven days of BMP4 treatment. However, as we increase the proliferation rate of PCs relative to GSCs (increase 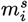) we observe two different possibilities depending on the compartment sensitivity (*φ*). If the compartment sensitivity is relatively low then PCs divide rapidly and can divide a number of times (around 10) before becoming TDCs, therefore the tumor size can increase over a short period of time. On the other hand, as we increase compartment sensitivity (*φ*) we observe that the tumor size after seven days of BMP4 treatment decreases, even for high 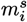. In this case the PCs rapidly become TDCs either straight away or after only a few divisions. Taken together these results show that there are two key parameters that determine the outcome of BMP4 exposure alone on tumor growth rate: i) the proliferative capacity of induced PCs (controlled by *φ*) and ii) the relative proliferation rate of PCs to GSCs 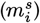. If PCs have a large proliferative capacity and proliferate faster than GSCs, BMP4 exposure can lead to an increase in size after seven days, while if PCs have a small proliferative capacity or proliferate slower than GSCs then BMP4 treatment will always lead to a decrease in size after seven days.

### 3.3. Continuous delivery of BMP4 is more effective than a single dose

Using our model, we simulate the effect of alterative BMP4 delivery schedules. Here we consider: A single dose at the time of resection (prior to RT), and a continuous dose from resection until the end of RT. In both cases the same total dose of BMP4 is given so the comparison is fair. A single dose could feasibly be delivered directly into the tumor bed at resection, whereas continuous dosing could be achieved via convection-enhanced delivery, as described in (37).

To compare over a wide range of parameter values that we expect to see across patients, we perform a large virtual clinical trial of 1000 virtual patients, each with a random set of parameters generated using Latin hypercube sampling. Each parameter is assumed to have a uniform distribution, the upper and lower bound for each parameter given in Table 2. The intentionally wide parameter ranges are chosen to attempt to capture the full spectrum of possible patient heterogeneity. Each virtual patient is then simulated undergoing three treatment arms: i) CTRL - resection followed by RT. ii) BMP4 single - a single dose of BMP4 at the time of resection followed by RT. iii) BMP4 continuos - a continuous dose of BMP4 from resection until the end of RT.

**Table 2:** Table of parameter values used for the large virtual trial of 1000 virtual patients, shown in Figure 4. When the value is fixed its value is given, if it is sampled from a uniform distribution then the lower and upper bounds are given. Dimensionless units are indicated with units of 1.

| Parameter | Meaning | Range/Value | Units | Reference |
| --- | --- | --- | --- | --- |
| $\delta_s$ | Death rate of GSCs | [0.0001, 0.002] | 1/year | Estimated |
| $\delta_i^s$ | Death rate of PCs relative to GSCs ($\delta_i = \delta_i^s \delta_s$) | [1, 20] | 1 | Estimated |
| $\delta_n^s$ | Death rate of TDCs relative to PCs ($\delta_n = \delta_n^s \delta_i$) | [1, 10] | 1 | Estimated |
| $\delta_m$ | Death rate of AMSCs | 0.5 | 1/year | (33) |
| $\delta_B$ | Decay rate of BMP4 | 0.5 | 1/year | (33) |
| $u_B$ | Uptake rate of BMP4 | [0.1, 1] | 1/year | Estimated |
| $C$ | Release rate of BMP4 from AMSCs | [0.1, 1] | 1/year | Estimated |
| $n$ | Proliferative capacity of PCs | 10 | 1 | (19) |
| $m_s$ | Proliferation rate of GSCs | [10, 90] | 1/year | Estimated |
| $m_i^s$ | Proliferation rate of PCs relative to GSCs ($m_i = m_i^s m_s$) | [1, 5] | 1/year | Estimated |
| $\psi$ | GSC self-renewal sensitivity | [0, 0.2] | 1 | Table 1 |
| $\varphi$ | GSC compartment sensitivity | [0, 0.1] | 1 | Table 1 |
| $\alpha^s$ | Radiosensitivity based on proliferation rate ($\alpha = \alpha^s m_s$) | [0.001, 0.01] | 1/Gy | (52) |
| $\beta$ | Quadratic component of RT cell killing | $\alpha/10$ | 1/Gy | (51,52) |
| $\eta$ | Difference in radiosensitivity between GSCs and PCs | [0.001, 1] | 1 | (19) |
| $\mu$ | Difference in radiosensitivity between PCs and TDCs | [0.2, 1] | 1 | (19) |
| $P_s^{\max}$ | Maximum rate of GSC self-renewal | [0.5, 0.6] | 1 | (28,58) |
| $P_s^{\min}$ | Minimum rate of GSC self-renewal | [0.1, 0.3] | 1 | (28,58) |

Figure 4 (A) shows the results for the top 100 virtual patients that had the largest improvement in survival time from BMP4 (and virtual patient 230 is added to show a poor responder). Over this large heterogenous population we found that for all patients, treating with continuous BMP4 delivery (red) was always equal to or superior than a single dose (blue). This is primarily due to the short half-life of BMP4, which means that a single dose is often not sufficient to maintain the GSC self-renewal rate *P*_*s*_ at a low level during the entirety of RT. This can be seen in the specific examples of virtual patients 900 and 820 Figure 4 (B-C). Both these patients show good response to BMP4, but when only a single dose is given the GSC fraction increases towards the end of RT when BMP4 has decayed. When BMP4 is delivered continuously, more is present during the final weeks of RT and thus the fraction of GSCs is kept reduced for the entire RT cycle. One of the virtual patients that showed poor response to BMP4 was patient 230, shown in Figure 4 (D). We further investigate the reasons for this heterogeneity in response in the following section.

**Figure 4:**
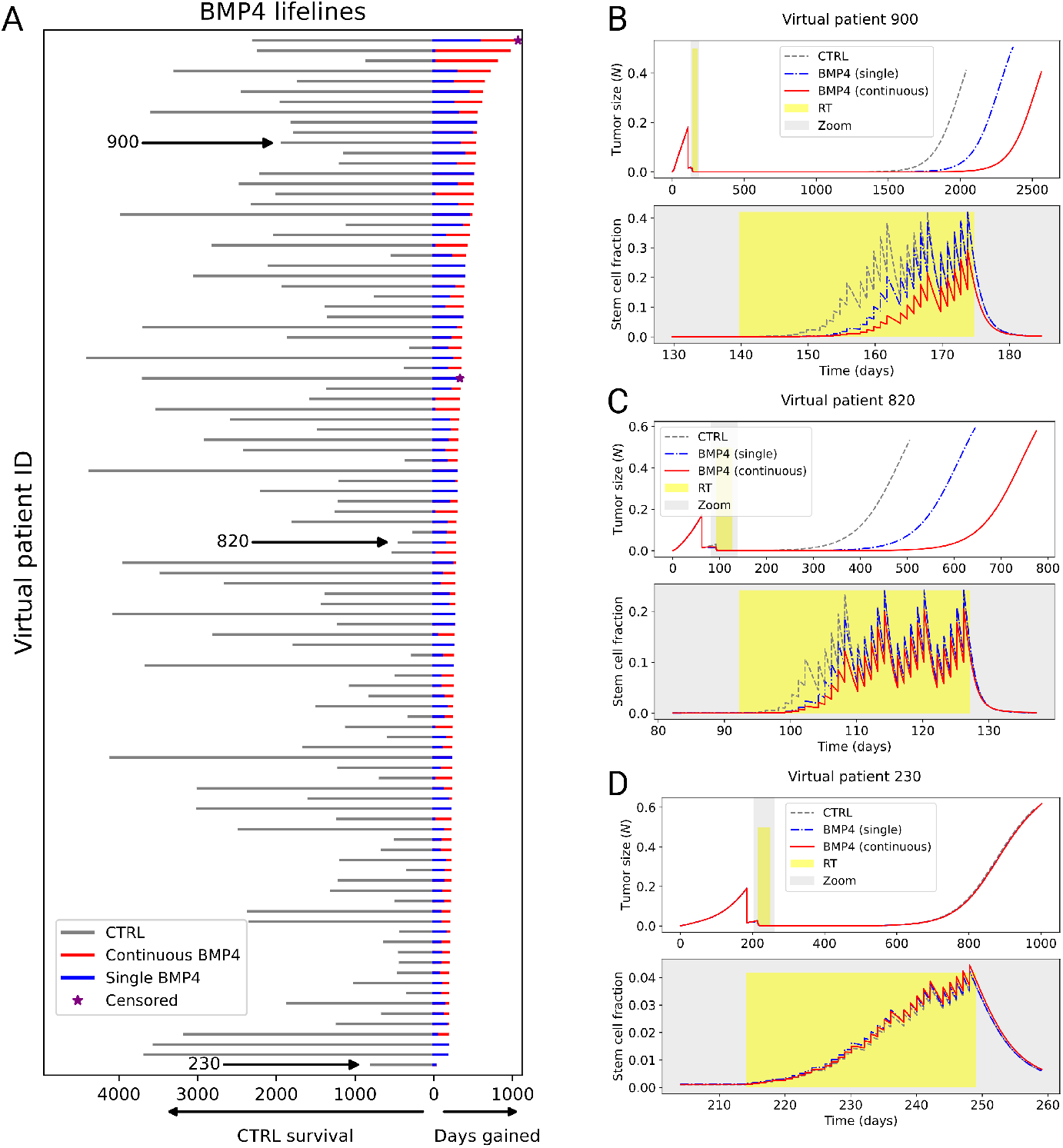
Continuous BMP4 treatment is more effective than single dose BMP4 and CTRL over a wide range of virtual patients. A) Lifelines plot for top-responding 100 out of 1000 virtual patients (and patient 230), showing survival time under CTRL in grey (to left) and the added days gained from BMP4 therapy (to right): single (blue) and continuous (red), ordered by days gained due to BMP4 therapy. Across all virtual patients days gained from continuous BMP4 was always greater than or equal to single BMP4; if no red line can be seen then single and continuous resulted in the same survival. In this virtual trial a patient is censored if they survive to the maximum simulation time, represented by the purple star. B)-D) Example tumor trajectories for three virtual patients, showing CTRL (grey), BMP4 single (blue) and BMP4 continuous (red). For each patient the top plot shows the total tumor size for the full simulation while the bottom plot zooms in around the time of RT (grey box) and shows the fraction of GSCs during this period. Virtual patient 900 and 820 both respond well to BMP4, particularly continuous delivery, while 230 has little response to BMP4.

### 3.4. Global sensitivity analysis identifies key parameters for successful BMP4 treatment

Given the large heterogeneity in response to BMP4 therapy observed in Section 3.3, we aim to identify the key parameters that determine its effectiveness. To determine the most important patient parameters, i.e., the ones that resulted in the largest improvement in survival between CTRL and BMP4 (continuous) arms we perform a linear regression global sensitivity analysis (GSA) on the model using the diverse 1,000 virtual patient cohort previously simulated in Section 3.3. The outcome metric (response variable) we compare is days gained fold change (DGFC). that is

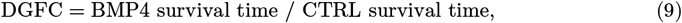

where survival time is the time from detection to death. Therefore, a DGFC score of less than 1 would indicate worse survival with BMP4 compared to CTRL, while a doubling in survival time with BMP4 would result in a DGFC score of 2. The explanatory variables of the linear regression model are the model parameters. Thus, the linear regression model is given by

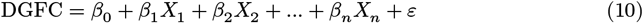

where *X*_*i*_ are the model parameters and *ε* is the error term. The coefficients *β*_*i*_ give the effect of each parameter on the DGFC. The larger the absolute value of the coefficient *β*_*i*_ the more effect it has on the resulting DGFC score. The sign of the coefficient dictates the direction of the effect. In order to make a fair comparison between the coefficients, we divide the parameters by their standard deviation, which gives us a measure of the relative importance of each parameter on the DGFC score. The results of this analysis are shown in Figure 5 (A). There are seven model parameters for which the 95% confidence interval of the standardizes slope (*β*_*i*_/ sd(*X*_*i*_)) does not include zero. Unsurprisingly, the two most important parameters are identified as *ψ* (GSC self-renewal sensitivity to BMP4) and *η* (the difference in radiosensitivity between GSCs and PCs). Also identified as important are *C* (the release rate of BMP4), *m*_*s*_ (the proliferation rate of GSCs), *α*^*s*^ (the relationship between proliferation rate and radiosensitivity), and 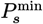 and 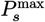 (the minimum and maximum self-renewal rates of GSCs). Of particular interest is the proliferation rate of GSCs, *m*_*s*_, as this could be potentially measured from a biopsy and selected for in a patient cohort. To isolate the effect of each parameter on DGFC, we used partial regression plots for the six most influential parameters. For each parameter, we first regressed DGFC on all other parameters and extracted the residuals, representing the variation in DGFC not explained by that parameter. Similarly, we regressed the parameter of interest on the remaining parameters and took the residuals, capturing its unique, independent variation. Plotting these residuals against each other (Figure 5 (C)) visualizes the parameter’s independent contribution to DGFC, with the slope corresponding to its regression coefficient.

**Figure 5:**
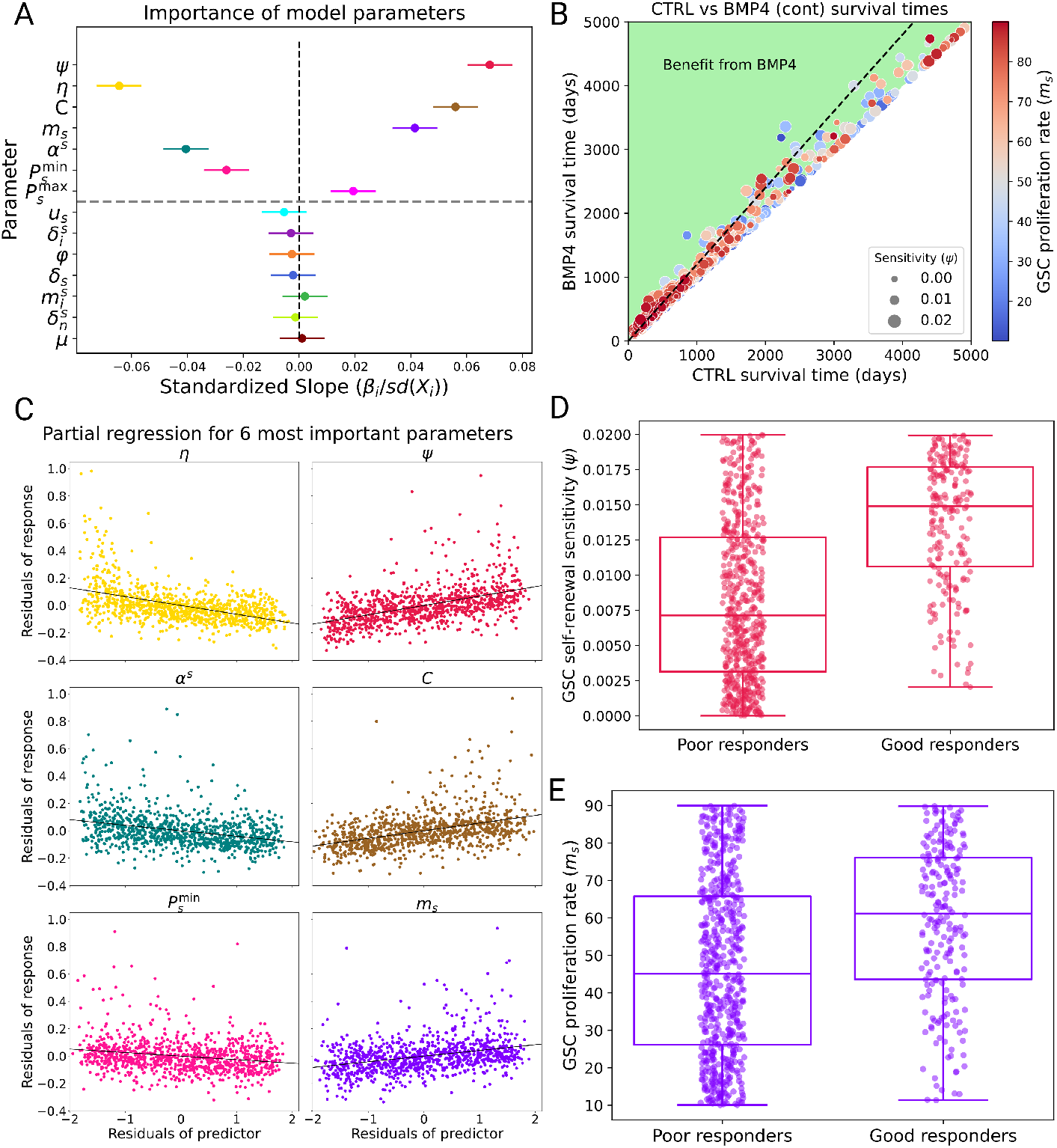
Global sensitivity analysis (GSA) identifies key parameters for successful BMP4 treatment. A) Standardized slope plot for all parameters sampled in the GSA. The dot represents the mean value of the coefficient (*β*_*i*_) and the bar the 95% confidence interval. If a parameter’s 95% confidence interval does not contain 0 (above grey dashed line), it can be expected to have a significant effect on DGFC. B) Plot visualizing the benefit of BMP4 considering both *m*_*s*_ (circle color) and *ψ* (circle diameter), the black dashed line represents a DGFC score of 1.2. C) Partial regression plots for the six most important model parameters. D)-E) We split the virtual patients into good responders (DGFC > 1.2) and poor responders (DGFC < 1.1). The virtual patients in the good responders group have on average higher *m*_*s*_ and *ψ* when compared to the poor responders.

To further investigate how GSC self-renewal sensitivity (*ψ*) and GSC proliferation rate (*m*_*s*_) affect the DGFC score, we consider two groups retrospectively, good responders (those who have a DGFC score of more than 1.2) and poor responders (those with a DGFC score of less than 1.1). We plot the distribution of *ψ* and *m*_*s*_ for these two groups in Figure 5 (D-E). There is a statistically significant difference in the *ψ* and *m*_*s*_ parameter distributions between the good and non-responders (independent two-sample t-test both *p* < 0.001), with the good responders on average having a higher *ψ* and *m*_*s*_ than the poor responders. Overall, these results verify that it is vital that potential clinical trial participants have some GSC self-renewal sensitivity (*ψ*) to BMP4 as well as a difference in radiosensitivity between GSCs and PCs (*η*), as without these BMP4 treatment cannot be effective. In addition, we also identify several other key model parameters that are associated with increased days gained under BMP4 treatment and, in particular, we suggest that patients with higher GSC proliferation rates (*m*_*s*_) may benefit more from BMP4 therapy.

### 3.5. Virtual clinical trials stratified by proliferation rate

Our goal was to investigate several patient selection criteria to identify potential patient cohorts that will observe the maximal benefit of an early phase clinical trial for a potential BMP4 therapy. The GSA in Section 3.4 identified parameters that significantly affect the DGFC score between CTRL and BMP4 arms. Previous studies have shown that proliferation rates of GBM tumor vary significantly between patients (44,59) and we have shown heterogeneity within GSC self-renewal sensitivity (*ψ*) among our twelve patient-derived cell lines (Table 1). As such, we consider patient stratification based on GSC self-renewal sensitivity *ψ* and GSC proliferation rate *m*_*s*_.

We create a total of five patient cohorts based on GSC proliferation rate *m*_*s*_ and GSC self-renewal sensitivity *ψ*, by splitting the full range of *m*_*s*_ and *ψ* given in Table 2 into the bottom (yellow) and top (blue) 50% of the parameter range, i.e., the parameter range for *m*_*s*_ [10, 90] is split into [10, 50] (yellow) and [50, 90] (blue), as well as considering a non-stratified group (green) that covers the full range. The same is done for *ψ*. This creates a total of five groups summarized in Figure 6 (A). Figure 6 (B) shows how this gives rise to three distinct distributions of *m*_*s*_ and *ψ* between these groups. All other patient parameters are again sampled from the uniform distributions given in Table 2.

**Figure 6:**
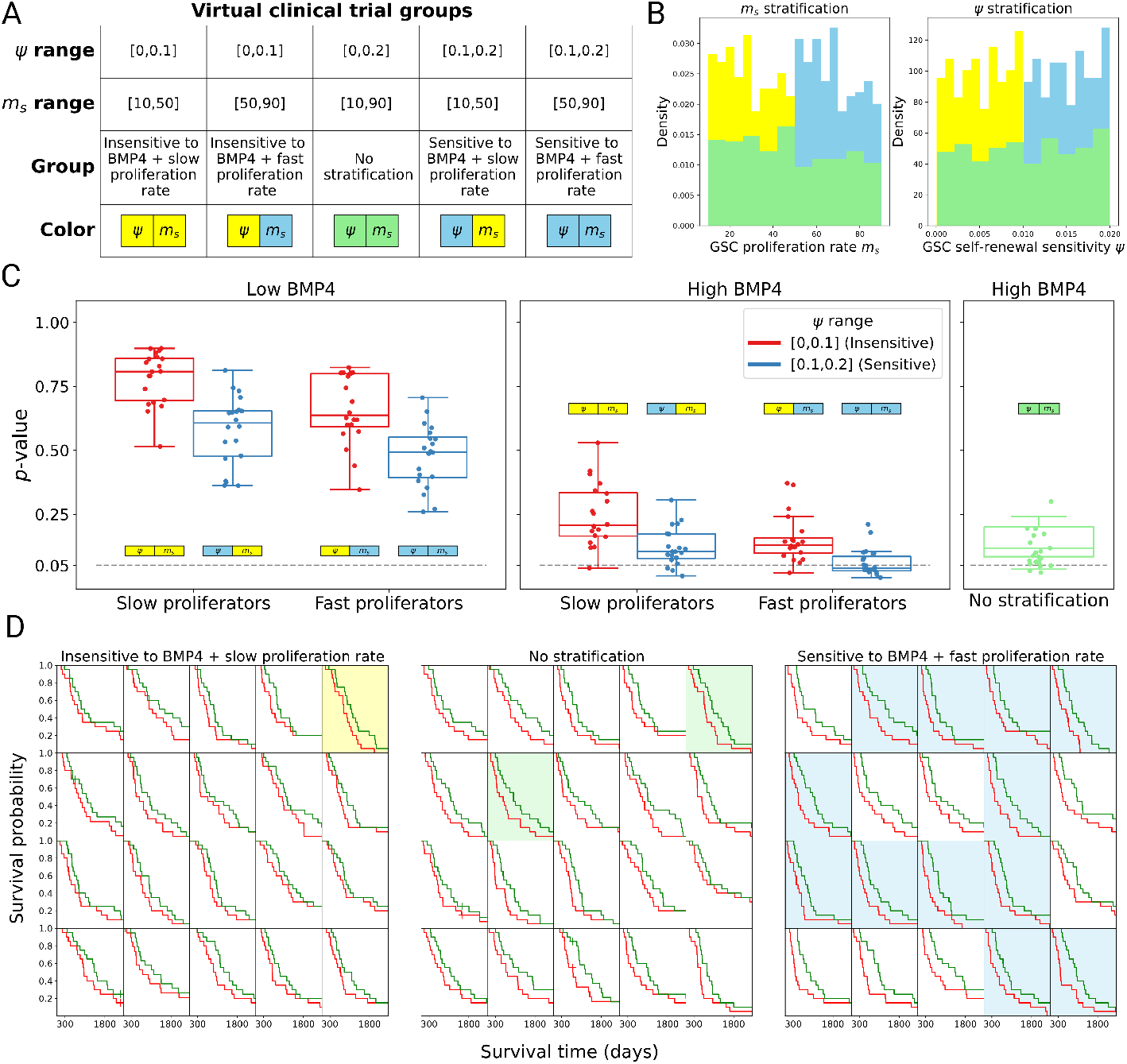
Stratified virtual clinical trials show the importance of patient selection. A) Table showing the parameter range for the five groups (four stratified and one non-stratified) on which we perform a series of virtual clinical trials B) Histogram showing how *m*_*s*_ and *ψ* are stratified. The bottom 50% group is in yellow, the top 50% group is in blue and the non-stratified group is in green. C) Summary of fraction of successful virtual trials (*p* < 0.05) across all cohorts considered. The grey dashed line indicates the significance level *p* = 0.05. D) Simulated Kaplan-Meier curves from the virtual trials for groups: insensitive and slow proliferation rate, no stratification and sensitive and fast proliferation (high BMP4), CTRL is plotted in red and BMP4 (continuous) is plotted in green.

For each stratified patient cohort, we perform a set of 20 virtual clinical trials, each trial containing 20 virtual patients. The number of virtual patients in each trial is chosen to roughly represent an early phase clinical trial. For the BMP4 arm, we perform these trials for two total concentrations of BMP4-AMSCs (low and high). For direct comparison, the CTRL arm consists of exactly the same virtual patients (same parameters) as the BMP4 arm. By performing a set of virtual clinical trials for each cohort we can estimate the probability of observing a successful trial, defined as the proportion of trials that have a statistically significant increase in survival time with BMP4 treatment (continuous) compared to CTRL (*p* < 0.05).

A summary of the results of these trials is shown in Figure 6 (C). If total BMP4-AMSCs delivered is too low, then as may be expected, we do not observe any significant difference between the BMP4 and CTRL trials. As we increase the total amount of BMP4-AMSCs delivered it becomes important to consider the patient cohort. First, when the mean GSC proliferation rate (*m*_*s*_) and GSC self-renewal sensitivity (*ψ*) is low, even if higher total doses of BMP4-AMSCs are achieved we are still unlikely to observe a statistically significant increase in survival time across a small virtual trial (1/20 successful). Second, if we consider the non-stratified cohort the large patient heterogeneity means we rarely see a robust signal for BMP4 (2/20 successful). Finally, if we instead consider the cohort with the highest mean proliferation rate and GSC self-renewal sensitivity, then we observe a higher fraction of successful trials (12/20). The full set of twenty virtual trials for each of these cohorts (under high BMP4) are shown in Figure 6 (D). All virtual trials show improved response under BMP4 treatment (green) compared to CTRL (red). However, groups 1 and 3 both contain a number of virtual patients with tumor characteristics that are associated with poor response to BMP4, therefore diluting the overall observed benefit of BMP4, resulting in many fewer trials identifying BMP4 as a potential treatment. These results highlight the importance of patient selection in early-phase clinical trials for BMP4-AMSC therapy.

## 4. Discussion

GSCs represent a significant barrier for successful treatment of GBM. They are both highly resistant to standard of care therapies (radiotherapy and chemotherapy), and have the unique capability to repopulate the tumor after treatment, even if just a small number survive. As such, if we are to improve patient outcomes for GBM it is vital that we find ways to target GSCs. BMP4 has shown some promise as a potential targeted therapy for GSCs, inducing differentiation of GSCs - increasing radiosensitivity and reducing overall tumor burden in *in vitro* and in animal studies (32,33), and recently has been shown to be safe in a phase I clinical trial (37). However, translating this into a successful clinical therapy faces many challenges. To begin addressing these challenges, we have developed a mathematical model for GSC proliferation and differentiation that incorporates the effects of BMP4 therapy as well as radiotherapy. Importantly, our model incorporates the differential radiosensitivity between GSCs, PCs and TDCs as well as the effects of BMP4 on GSC self-renewal and the proliferative capacity of induced PCs. This mathematical model can be used to simulate a wide range of patient profiles and treatment schedules, allowing us to thoroughly analyze the potential of BMP4 therapy *in silico*.

Despite its potential as a therapy, relatively few mathematical models have focused on the potential impact of such an agent on tumor growth (18,28,58,60). Previous models have focused on general differentiation therapy agents and show results consistent with our findings, that differentiation therapy could be beneficial for tumor control (28,58). To the best of our knowledge, Turner *et al*. (18) is the only previous mathematical model that has specifically considered BMP4 as a potential differentiation therapy for GBM. However, their model is derived from a stochastic birth-death process, and they did not attempt to quantify sensitivity to BMP4, explore multiple treatment schedules or identify parameters that are important for successful BMP4 therapy. Interestingly, their model also predicted that BMP4 could lead to both an increase and decrease in size after treatment.

To quantify the effects of BMP4, we performed both a RT assay and a proliferation assay on twelve patient-derived GSC lines. The RT assay showed that, apart from a subset of non-responders, BMP4 increased the sensitivity to RT of GSC lines (Figure 2). The proliferation assay showed that GSC lines could both increase or decrease in size, relative to control conditions, after seven days of exposure to BMP4. We demonstrate that, in order for our model to reproduce the results of both the RT assay and the proliferation assay, we must introduce an effect of BMP4 on GSC self-renewal (*ψ*) and the proliferative capacity of induced PCs (*φ*). By introducing these model parameters and fitting the model to the experimental data we are able to estimate the sensitivity of each cell line to BMP4, both in terms of GSC self-renewal sensitivity (*ψ*) and the proliferative capacity of PCs (*φ*). Having quantitative model based measures of sensitivity to BMP4 is vital for understanding heterogeneity in treatment response.

Using our mathematical model we simulated a cohort of 1000 virtual patients. Due to limited information on some of the model parameters we do not attempt to draw them from parametrized distributions, but instead consider a wide range on each parameter and sample from a uniform distribution (using Latin hypercube sampling). This allows us to explore the full range of potential patient profiles. We simulated a virtual clinical trial comparing resection and RT (CTRL) with a single dose of BMP4-AMSCs at the time of resection followed by RT and a continuous dose of BMP4-AMSCs from resection until the end of RT. This showed that, over the wide range of parameter space sampled, the continuous delivery of BMP4-AMSCs was always more effective than a single dose. This is primarily due to the short half-life of BMP4 and the synergy it has with RT. If BMP4 is still present during RT it is able to reduce the enrichment of GSCs, this results in greater cell killing and delayed time to recurrence.

To identify the key model parameters that resulted in a benefit from BMP4-AMSC therapy we performed a global sensitivity analysis (GSA) on the model, utilizing the same 1000 virtual patients. Linear regression analysis identified seven key parameters that had a significant effect on improving survival time with BMP4-AMSC therapy (continuous delivery): GSC self-renewal sensitivity (*ψ*), the difference in radiosensitivity between GSCs and PCs (*η*), the release rate of BMP4 from AMSCs (*C*), the proliferation rate of GSCs (*m*_*s*_) and the maximum and minimum probabilities of self-renewal (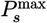 and 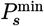). Many of the parameters identified are intuitive as they directly relate to BMP4 treatment. Of particular interest, the GSA identified the GSC proliferation rate (*m*_*s*_) as an important parameter for successful BMP4-AMSC therapy. This agrees with previous biological studies that have suggested the effects of BMP4 might be more profound in tumors with high proliferation rates (37). Additionally, we believe proliferation rate has the potential for patient stratification in clinical trials, since it shows significant variation between patients and is measurable either from a biopsy or inferred from MRI.

Several challenges remain with the development of BMP4-AMSC therapy for GBM. Chief among them is identifying why some cell lines respond to BMP4 while others do not. Previous single cell RNA-seq studies have identified Olig1/2 expression as vital for sensitivity to BMP4 between cell lines (34). In the recent phase I clinical trial it was found that BMPR1B expression had a significant association with patient outcome (37). These have the potential to be used as biomarkers to predict BMP4 sensitivity, however further work is needed to characterize the mechanisms that underlie BMP4 sensitivity/resistance in GSCs.

All mathematical models rely on simplifying assumptions, and ours is no exception; we do not claim to capture the full complexity of a GBM tumor or its response to BMP4-AMSC therapy, which itself is not yet fully understood. Our model, while mechanistically detailed in capturing the hierarchy of GSCs, PCs and TDCs, simplifies the spatial heterogeneity of GBM and the tumor microenvironment. It assumes uniform BMP4 delivery and does not incorporate feedback mechanisms or immune interactions. Future modelling efforts could focus on addressing these limitations, particularly, due to evidence that BMP4-AMSCs can migrate towards GSCs (35) and that GSC create an immunosuppressive microenvironment (61).

Finally, we showed how we can use our virtual tumor digital twin to inform patient selection for successful clinical trial design. If outcomes for GBM patients are to be improved there is an urgent need for new therapies, particularly ones that target GSCs. But developing new treatments is challenging and designing clinical trials is notoriously difficult and fraught with challenges, including patient selection and controlling for patient heterogeneity. We believe that by incorporating a mathematical model digital twin right from the beginning it can help guide this process, identifying patient tumor characteristics that drive maximal benefit (and those that are not important or do not need to be controlled for), and allowing us to identify potentially novel tumor biomarkers that can be used for patient selection during clinical trials.

## Acknowledgments

This publication was made possible through the support of the Distinguished Mayo Clinic Investigator Award (A.Q.-H.), Monica Flynn Jacoby Endowed Chair (A.Q.-H.), the William J. and Charles H. Mayo Professorship (A.Q.-H.), the Florida Department of Health Cancer Research Chair Fund (A.Q.-H.), the Richard and Lauralee Uihlein Neuro-oncology Research Fund (A.Q.-H.), the Jacquie Lorraine Goldman Fund for a Brain Tissue Bank (A.Q.-H.), the Florida State Funds for the Casey DeSantis Cancer Research Program (A. Q-H) and Current NIH Funding (A.Q-H): R01NS129671, R01CA282451, R01CA284268, U54CA27504 (K.R.S.), and U01CA250481 (K.R.S.). The cell lines (GBM1A & GBM965) utilized in this investigation were established from invaluable patient-derived specimens collected through the Mayo Clinic Florida Neurosurgery BRIDGE Biobank, a central nervous system tumor bioresource that facilitates translational research and advances scientific discovery on a global scale.

