## Supplementary Information for "Virtual Clinical Trials of BMP4 Differentiation Therapy: Digital Twins to Aid Successful Glioblastoma Trial Design"

### S.1 Parameter fitting

To parametrize the key model parameters of BMP4 sensitivity $\psi$ and $\varphi$ we utilize our model to simulate both the RT assay and the proliferation assay, we then perform a joint fit to minimize the overall error between the simulated and experimental data across both simulated assays. In all simulations we use the measured cell line doubling times to set the GSCs proliferation rate $m_s$, and the other parameters used are fixed across all cell lines according to the values in Table 1 (below). The final fitted parameters for each cell line are shown in Figure 1.

| Parameter | Meaning | Value | Units |
|---|---|---|---|
| $\delta_s$ | death rate of GSCs | 0.001 | 1/year |
| $\delta_i$ | death rate of PCs | 0.01 | 1/year |
| $\delta_n$ | death rate of TDCs | 0.1 | 1/year |
| $n$ | Proliferative capacity of PCs | 10 | 1 |
| $m_s$ | Proliferation rate of GSCs | Taken from doubling time of cell line | 1/year |
| $m_i^s$ | Proliferation rate of PCs relative to GSCs ($m_i = m_i^s m_s$) | 2 | 1 |
| $\eta$ | Difference in radiosensitivity between GSCs and PCs | 0.1376 | 1 |
| $\mu$ | Difference in radiosensitivity between PCs and TDCs | 0.5 | 1 |
| $P_s^{\max}$ | Maximum rate of GSC self-renewal | 1 | 1 |
| $P_s^{\min}$ | Minimum rate of GSC self-renewal | 0 | 1 |

Table S1: Table of parameter values used for fitting the model to cell line data (RT and proliferation assay)

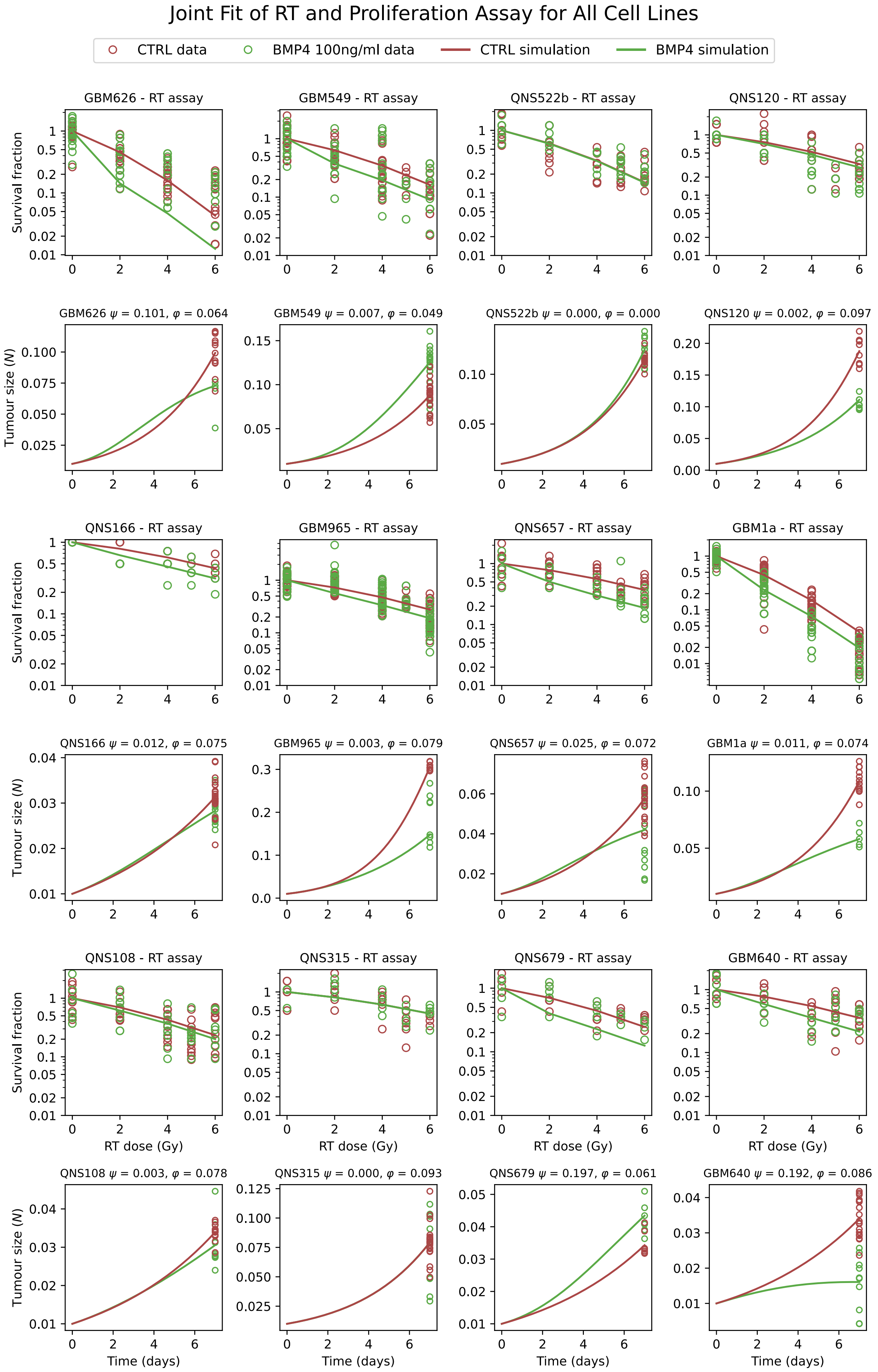


Figure S1: Results of the joint fit of both GSC self-renewal sensitivity ($\psi$) and compartment sensitivity ($\varphi$) for all cell lines

### S.2 Adipose derived mesenchymal stem cells as delivery mechanism for BMP4

The MSC were isolated from the lipoaspirates obtained from the consenting healthy donors after the approval from the Mayo Clinic Institutional Review Board. The fat tissue was collected and transferred to the laboratory and processed under sterile conditions and cultured in a GMP-compliant culture media (1). The isolated primary cell lines were maintained in GMP-compliant culture media and validated for MSC characteristics. The cells were sub-cultured upon 70% confluency and used for further experiments.

MSCs were transduced with a lentiviral vector engineered to express BMP4 (2), utilizing a multiplicity of infection (MOI) of 10 in the presence of 4 μg/mL polybrene in complete media. Transduction proceeded for 72 hours. Subsequently, transduced cells were isolated using a FACS Aria cytometer to enrich for transgene-expressing cells.

#### S.2S.1 BMP4-AMSC sensitivity

For, five of the cell lines (GBM1a, GBM965, GBM626, QNS120) we also performed the radiotherapy assay with BMP4-AMSCs. The results are plotted below. In general BMP4-AMSC behaves similarly to BMP4 alone, increasing radiosensitivity.

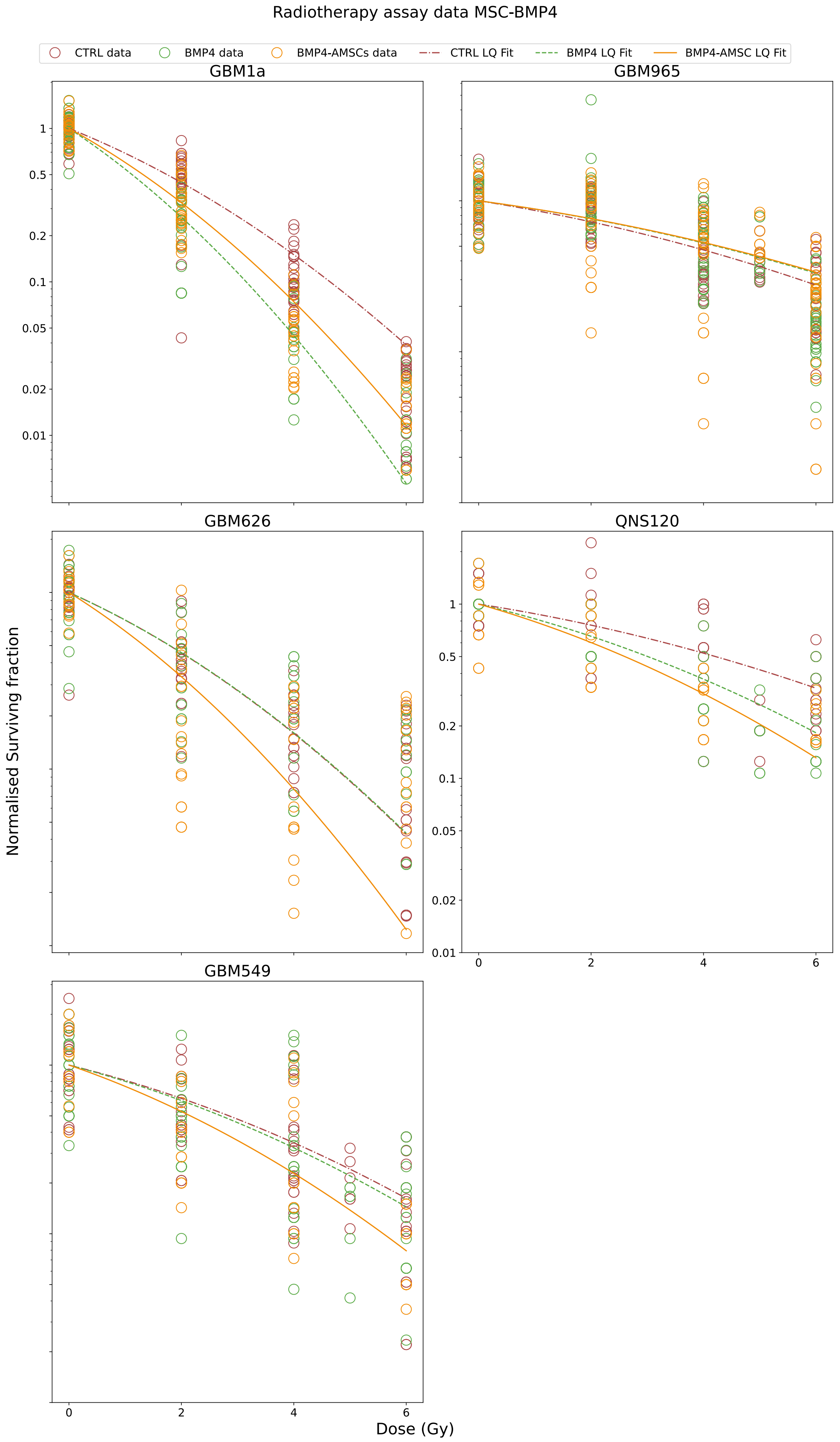


Figure S2: Radiotherapy assay results with CTRL (red, dashed), BMP4 (green, dash-dot) and BMP4-AMSC (orange). In all conditions th LQ model is fit soley for illustrative purposes

### S.3 Latin hypercube samples for 1000 patients virtual clinical trial

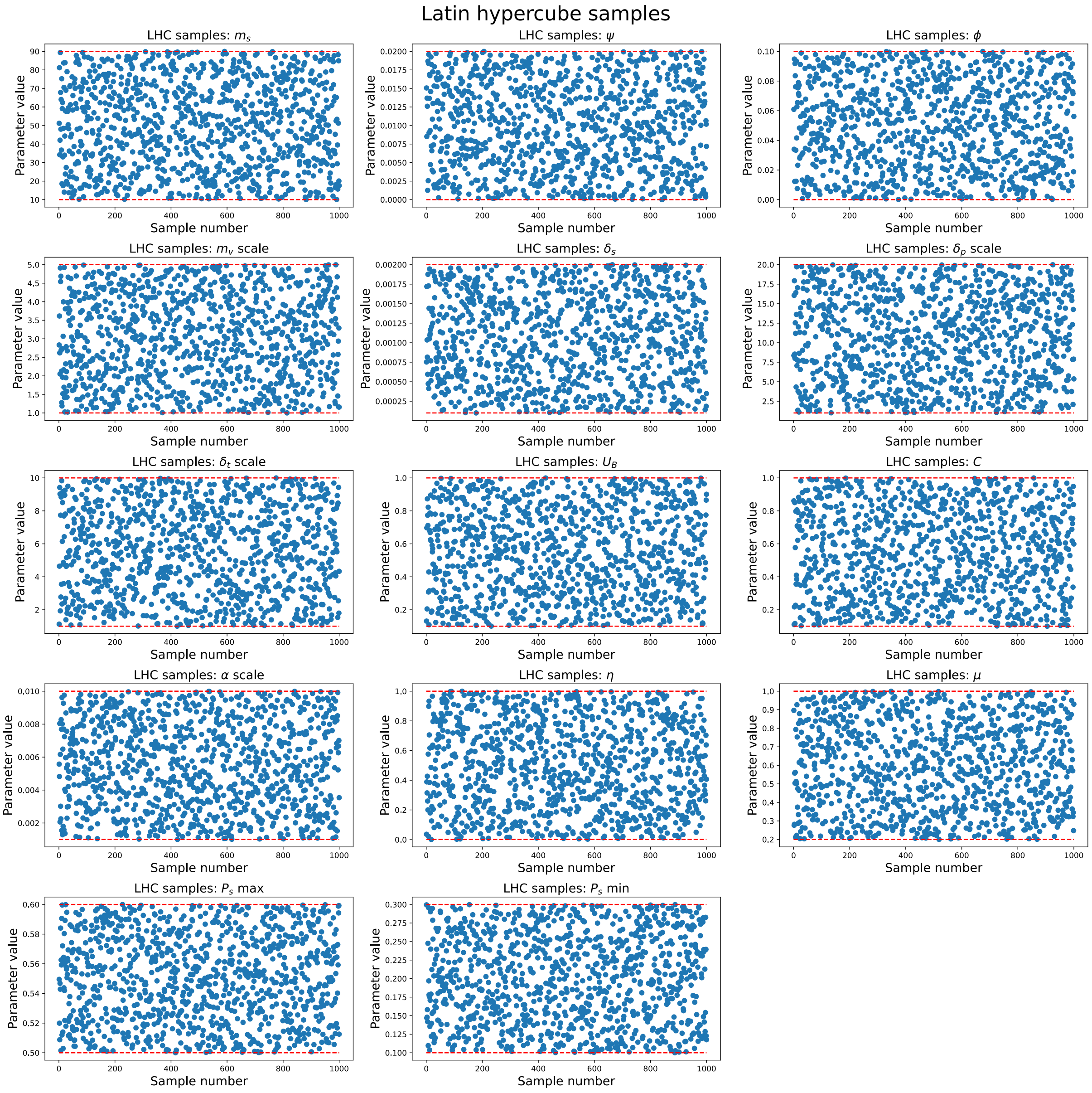


Figure S3: Latin hypercube samples for all model parameters used in large virtual clinical trial consisting of 1000 virtual patients.

### S.4 Relationship between parameter value and days gained

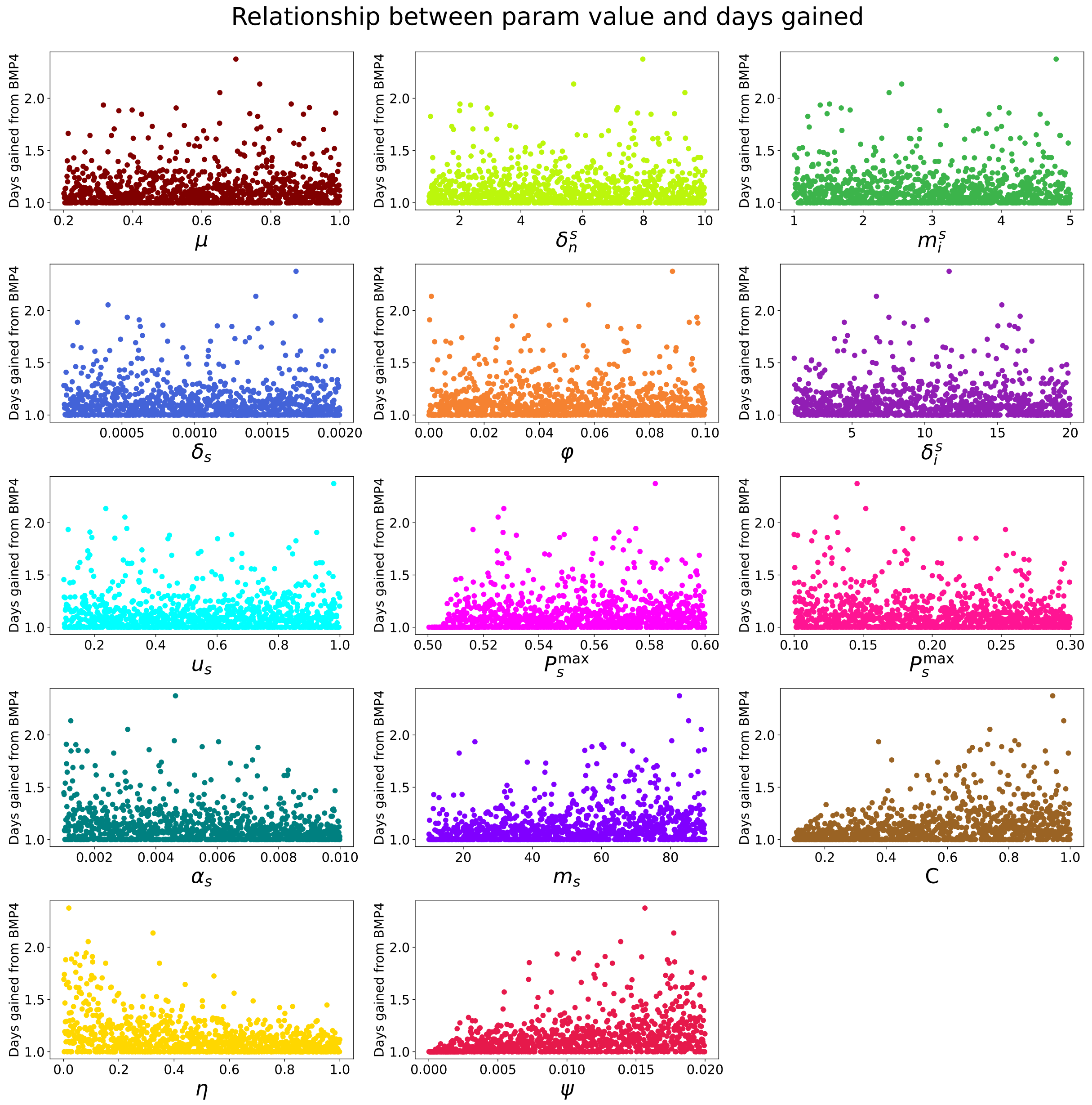


Figure S4: Direct relationship between parameter value and days gained for all model parameters sampled in the large virtual clinical trial of 1000 virtual patients.
